# Micronutrient supplements with iron promote disruptive protozoan and fungal communities in the developing infant gut

**DOI:** 10.1101/2021.07.06.451346

**Authors:** Ana Popovic, Celine Bourdon, Pauline W. Wang, David S. Guttman, Sajid Soofi, Zulfiqar A. Bhutta, Robert H. J. Bandsma, John Parkinson, Lisa G. Pell

## Abstract

Supplementation with micronutrients, including vitamins, iron and zinc, is a key strategy to alleviate child malnutrition. However, adverse events resulting in gastrointestinal disorders, largely associated with iron, has resulted in ongoing debate over their administration. To better understand their impact on gut microbiota, we analysed the bacterial, protozoal, fungal and helminth communities of stool samples collected from children that had previously been recruited to a cluster randomized controlled trial of micronutrient supplementation in Pakistan. We show that while bacterial diversity was reduced in supplemented children, vitamins and iron may promote colonization with distinct protozoa and mucormycetes, whereas the addition of zinc ameliorates this effect. In addition to supplements, residence in a rural versus urban setting is an important determinant of eukaryotic composition. We suggest that the risks and benefits of such interventions may be mediated in part through eukaryotic communities, in a manner dependent on setting.

## Introduction

Malnutrition is a global health crisis with 149 million children stunted and 45 million children wasted under the age of five years^1,2^. With increased vulnerability to infection, undernourished children are at elevated risk of death, not least from diarrheal diseases^3,4^. Previous studies have demonstrated the role of gut microbiota in malnutrition, with microbiome immaturity (bacterial communities that are underdeveloped with respect to age) representing a key factor in disease development^5,6^. Beyond bacterial communities, parasites such as hookworm, *Cryptosporidium* and *Entamoeba* have also been associated with severe diarrheal disease and intestinal malabsorption^7,8^. However, much less is known regarding the role of other, potentially commensal, eukaryotic gut microbes in undernutrition. Of particular interest is their ability to interact with and alter bacterial communities. For example, indole-producing gut bacteria were found to confer protection against *Cryptosporidium* infection, while deworming treatments targeting helminth endemic communities reduced abundance of protective Clostridiales^9,10^. Mouse studies further showed that helminths and protozoa influence bacterial communities by modulating the host immune system^9,11,12^. While the number of published gut microbiome studies have increased rapidly over the last decade, few have explored the composition of eukaryotic gut communities and their potential interactions with bacteria. Previously, we applied 18S rRNA and internal transcribed spacer (ITS) sequence surveys to systematically characterize eukaryotic microbiota in severely malnourished Malawian children, and identified a high prevalence of protozoa, including commensals and pathobionts^13^. We furthermore associated *Blastocystis* colonization with increased gut bacterial diversity.

Global health programs targeting vulnerable child populations include the use of micronutrient supplements, consisting of vitamins as well as essential minerals zinc and iron, that have been demonstrated to improve growth and reduce morbidity^14–16^. Such supplements are thought to address deficiencies that can impair immune responses to infectious pathogens and impact gut bacterial communities^17–20^. While beneficial, supplementation, especially with iron, may also promote unintended pathogen growth, particularly where the host is unable to restrict micronutrient bioavailability^21^. For example, it has been shown that surplus iron promotes the growth of enteropathogens and induces intestinal inflammation in infants^22,23^. Furthermore, while known to reduce the duration of childhood diarrheal episodes, zinc supplementation has been associated with increased duration of *Entamoeba histolytica* infections^24,25^.

In an attempt to understand the impact of micronutrient supplementation on the complex interactions between eukaryotic and bacterial microbiota in the maturing infant gut and health, we performed 18S rRNA and 16S rRNA amplicon surveys on stool samples obtained at 12 and 24 months of age from 80 children, previously recruited as part of a cluster randomized trial conducted in Pakistan. The trial was designed to investigate the impact of micronutrient powders (MNP) containing vitamins and iron with or without zinc on growth and morbidity, and has shown an excess of significant diarrheal and dysenteric episodes among children receiving MNPs^26^. Microbial profiles were analysed in the context of supplementation, nutritional status, age and place of residence (i.e., urban or rural) to reveal a complex landscape of associations with microbial diversity, as well as specific taxa.

## Results

### Description of cohort

A total of 80 children (160 paired stool samples at 12 and 24 months of age) from all three supplementation arms in the parent cRCT^26^ (control (n=24), MNP (n=29), and MNP with zinc (n=27)) conducted in Sindh, Pakistan were selected based on sample availability for inclusion in this study (Supplementary Fig. 1). The cohort includes children from both urban (Bilal colony) and rural (Matiari district) study sites (Fig. 1a). Children were stratified by weight-for-length z-scores (WLZ) at 24 months into a reference WLZ (WLZ >−1) or undernourished (WLZ < −2) group. Subject characteristics are summarized in Table 1. The WLZ growth trajectories of the children selected as the reference WLZ group approximately tracked the upper 50th percentile of the original cohort, while the undernourished group started around the lower 50th percentile and gradually dropped over time ending at the bottom 80^th^ percentile of the cohort (Fig. 1b). This drop in the WLZ of the undernourished children was driven by poor weight gain (Supplementary Fig. 2).

**Table 1.** Participant characteristics. Categorical values are presented as n (%), continuous variables show the mean and 95% confidence intervals. Premature birth was defined as gestational age < 37 months. Initiation of breastfeeding was reported for the period prior to recruitment into the study.

|  | **Undernourished**<br>(n=31) | **Reference WLZ**<br>(n=49) | **Total**<br>(n=80) |
| --- | --- | --- | --- |
| **Rural site, n(%)** | 25 (80.6%) | 28 (57.1%) | 53 (66.2%) |
| **Treatment arm, n(%)** |  |  |  |
| Control | 10 (32.3%) | 14 (28.6%) | 24 (30.0%) |
| MNP | 14 (45.2%) | 15 (30.6%) | 29 (36.2%) |
| MNP with zinc | 7 (22.6%) | 20 (40.8%) | 27 (33.8%) |
| Female, n(%) | 14 (45.2%) | 30 (61.2%) | 44 (55.0%) |
| Premature birth, n(%) | 6 (19.4%) | 10 (20.4%) | 16 (20.0%) |
| Initiated breastfeeding, n (%) | 31 (100.0%) | 47 (95.9%) | 78 (97.5%) |
| **Anthropometry, 12 mo** |  |  |  |
| Weight, Kg | 6.6 (6.3, 7.0) | 8.5 (8.2, 8.8) | 7.8 (7.5, 8.1) |
| Length, cm | 69.2 (67.7, 70.7) | 71.2 (70.4, 72.1) | 70.6 (69.9, 71.4) |
| Weight-for-length, z-score | -2.4 (-3.1, -1.7) | -0.0 (-0.3, 0.2) | -0.7 (-1.1, -0.4) |
| **Anthropometry, 24 mo** |  |  |  |
| Weight, Kg | 8.0 (7.6, 8.3) | 10.5 (10.2, 10.9) | 9.5 (9.2, 9.9) |
| Length, cm | 78.9 (77.3, 80.4) | 80.5 (79.5, 81.4) | 79.8 (79.0, 80.7) |
| Weight-for-length, z-score | -2.9 (-3.2, -2.7) | 0.2 (-0.1, 0.4) | -1.0 (-1.4, -0.7) |

**Fig. 1.**
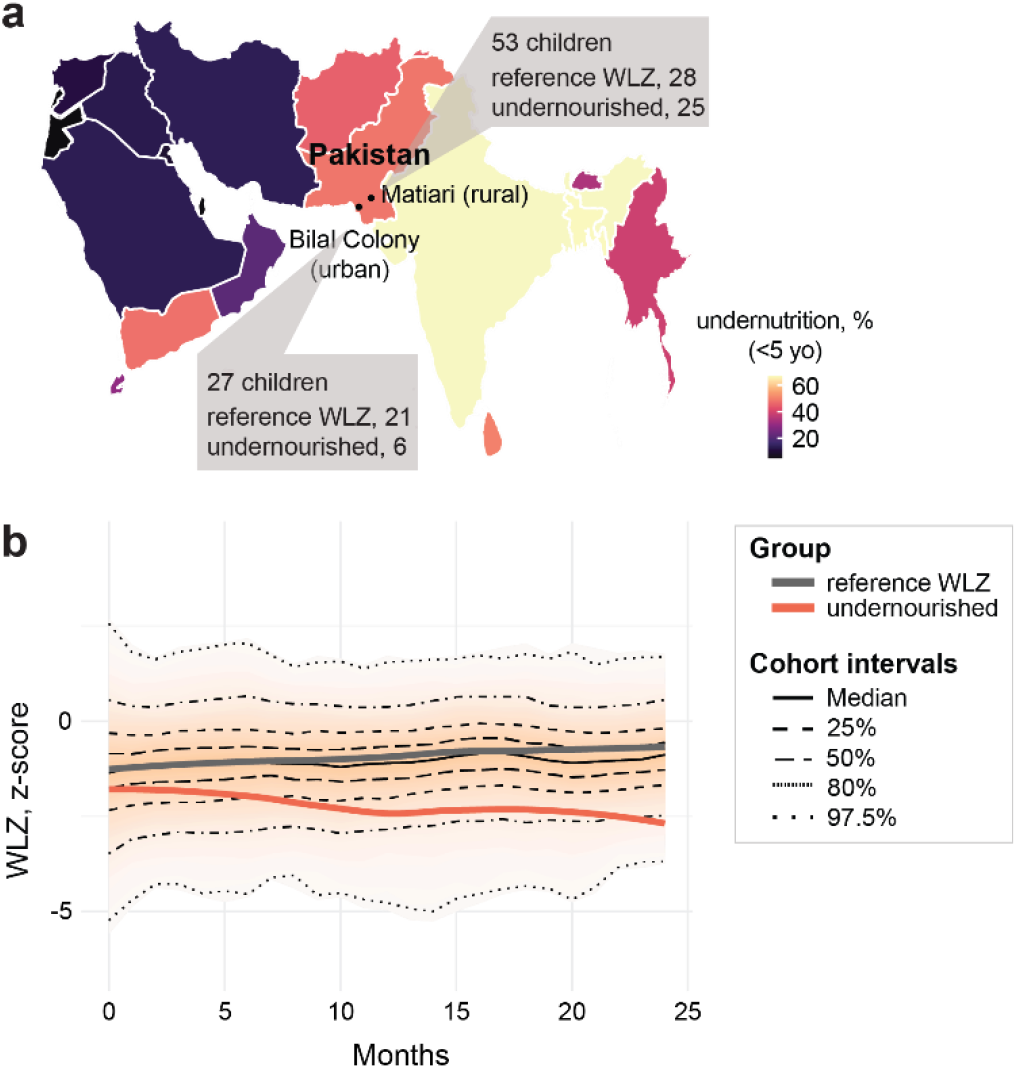
Participant characteristics. (a) Level of childhood undernutrition in Pakistan and the surrounding regions. Latest country data was retrieved from www.who.int/data/gho/indicator-metadata-registry/imr-details/27 on Feb 1, 2021. Urban and rural places of residence of the participants are indicated. (b) Weight-for-length z-scores of children recruited into clinical trial NCT00705445 during the first 24 months of life. Median and quantile values are shown, with medians for participants profiled in current study indicated by red (undernourished) and black (reference WLZ) lines.

**Fig. 2.**
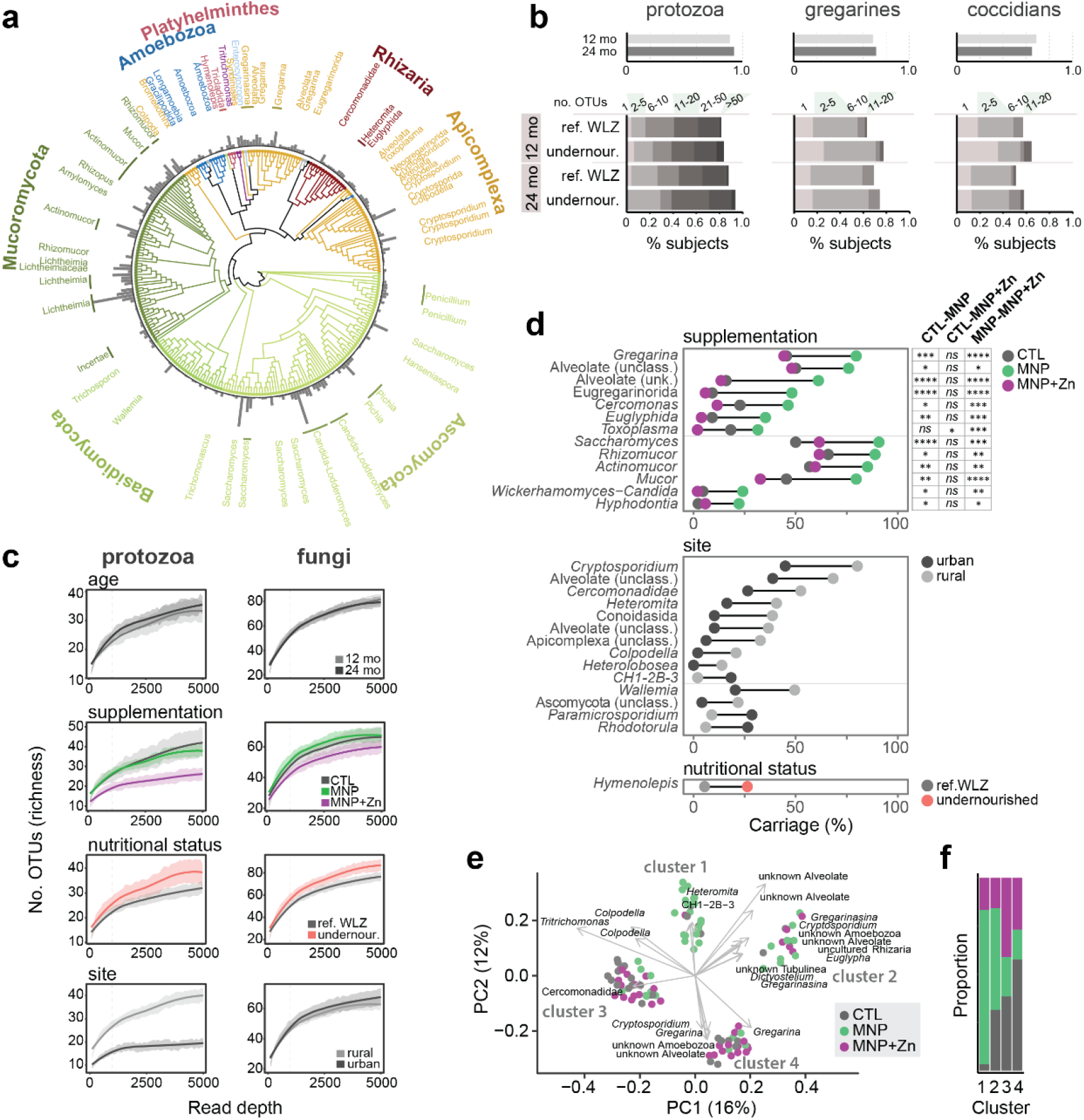
Eukaryotic communities in the gut are diverse and impacted by micronutrient supplementation and place of residence. (a) Phylogenetic tree representing eukaryotic taxa detected in children. Branches are coloured by phylum and bars represent the prevalences of OTUs in the cohort. Named organisms represent those detected in more than 5% of samples with a minimum of 100 reads. (b) Prevalences of protozoan (left), and specifically gregarine (middle) or coccidian (right) OTUs detected in children at 12 and 24 months of age. Prevalences are subdivided by nutritional group in bottom graphs, where shaded regions denote binned numbers of OTUs identified per sample. (c) Rarefaction curves comparing the mean protozoan and fungal species richness by age group, micronutrient supplementation, nutritional status and place of residence (site). Shaded regions represent standard error. Dashed lines denote the read depth at which significance was tested. (d) Carriage of eukaryotic taxa significantly associated with micronutrient supplementation, place of residence (site) or nutritional status. Results from Fisher’s pairwise tests among supplementation groups are indicated to the right. \**p* < 0.05, \*\**p* < 0.01, \*\*\**p* < 0.001. (e) Principal coordinate analysis of sample dissimilarities (n=106) based on protozoan composition, calculated using unweighted Unifrac scores. Samples are coloured by supplementation arm, and arrows indicate the direction of cosines of taxa significantly correlated with the first two principal components. Arrow lengths are scaled by the root square (r2) of the correlation. Identified clusters are numbered 1 though 4. (f) Proportions of samples from the respective supplementation arms within each protozoan community cluster.

### The developing infant gut is colonized by complex eukaryotic communities

We applied 18S rRNA amplicon sequencing to profile the eukaryotic communities in all 160 stool samples. We generated a total of 11,639,233 paired 18S rRNA amplicon sequence reads (median 70,642) of which 4,386,494 could be classified as a eukaryotic microbe (median 22,932; Supplementary Table 1). From these we identified a total of 859 eukaryotic OTUs (median 66; Supplementary Table 1), which included 438 protozoan, three helminth and 418 fungal OTUs (Fig. 2a). Fungi, dominated by Mucoromycota and Ascomycota, accounted for 71% of all reads. The most abundant were species in the *Candida-Lodderomyces* clade, *Saccharomyces*, and taxa increasingly associated with rare but fatal infections known as mucormycoses: *Rhizomucor*, *Actinomucor* and *Lichtheimia*. Alveolates accounted for 25% of reads, with *Gregarina*/*Gregarinasina* and *Cryptosporidium* as the most abundant (5% and 3%, respectively). Remaining reads were classified to numerous taxa, including known gut parasites such as *Enterocytozoon bieneusi*, *Pentatrichomonas hominis* and the tapeworm *Hymenolepis nana*, as well as uncharacterized alveolates, Amoebozoa and Cercozoa (Supplementary Table 2).

Protozoa were highly prevalent, with 89% of children colonized by at least one protozoan organism by 12 months of age, and 92% by 24 months of age (Fig. 2b). Carriage of multiple species was common in both the reference WLZ and undernourished groups, with on average 18 and 19 OTUs per child at each time point, and a maximum of 91. A high detection of gregarines, typically considered parasites of invertebrates, has not previously been reported in the human gut. In our cohort, gregarine sequences accounted for nearly 230,000 reads and were identified in 69% and 71% of children at 12 and 24 months of age (Fig. 2b).

### Micronutrient supplementation without zinc is associated with increased carriage of protozoa and mucormycetes

Protozoan microbiota were significantly associated with place of residence, micronutrient supplementation and/or nutritional status, but not age. Children residing in the rural study site had increased protozoan richness (number of OTUs) compared to those from the urban setting (β = 11, CI [5.3 – 16.6], *p* < 0.001) (Fig. 2c). Differences were attributed to higher carriage of predominantly alveolate taxa, particularly *Cryptosporidium* (Fisher’s exact, CI [2-11], *p* < 0.01, OR 4.9), species known to cause enteric symptoms (Fig. 2d). When stratifying by age group, only *Cryptosporidium* and two OTUs classified as unknown Conoidasida, with 93% sequence identity to *Cryptosporidium*, reached statistical significance at 24 months, with 2.4 and 9.6-fold higher carriage, respectively, in children from rural settings (Fisher’s exact, CI [2.5-29], *p* < 0.05, OR 8.1; CI [2-670], *p* < 0.05, OR 15.2).

While we observed trends in increased fungal and protozoan richness in the undernourished cohort (Fig. 2c), only the tapeworm *Hymenolepis nana* was detected with overall significantly higher frequency in undernourished children (Fisher’s exact, CI [2-23], *p* < 0.05, OR 6.2) (Fig. 2d). At 12 months, detections were only 2% and 3% in reference WLZ and undernourished children, respectively. However, by 24 months, carriage increased to 8% in reference WLZ and 43% in the undernourished group (*ns* after multiple testing correction). We also observed trends of increased carriage of *Cryptosporidium* and *Cryptosporida* (coccidians), represented by 46 OTUs in total, in undernourished children (74% versus 65% at 12 months and 71% versus 61% at 24 months; *ns*) (Fig. 2b). Furthermore, undernourished children receiving MNP with zinc had significantly fewer protozoan OTUs relative to undernourished children in the control and MNP arms (β = −15.19, CI [−29.27 – −1.12], *p* < 0.05), suggesting a possible inhibitory effect by the metal (Fig. 2c, Supplementary Fig. 3).

**Fig. 3.**
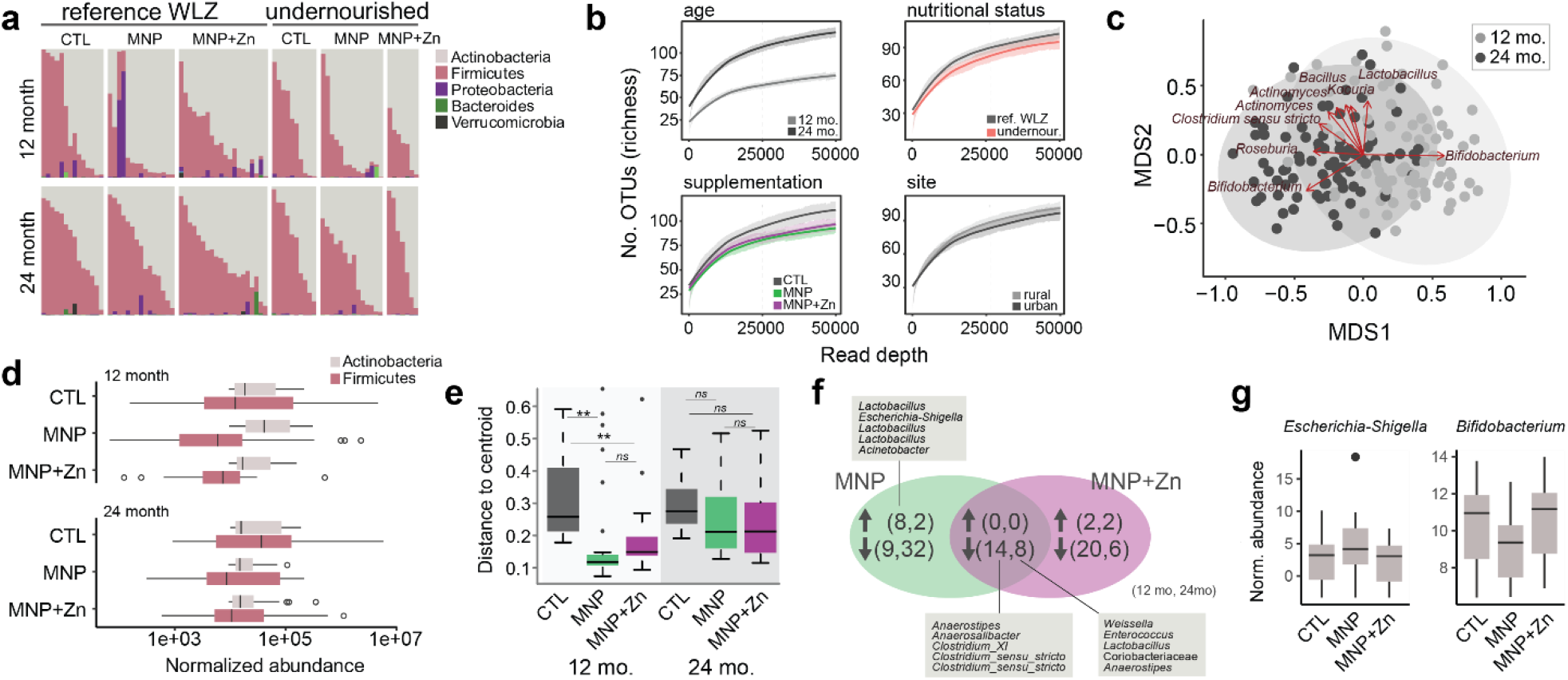
Bacterial microbiota change with age and supplementation. (a) Relative abundances of bacterial phyla in 12 (top) and 24 (bottom) month old children based on 16S data. Samples are sorted by the proportion of Firmicutes along the horizontal axis. (b) Rarefaction curves comparing mean species richness by age group, micronutrient supplementation, nutritional status and place of residence (study site). Shaded regions represent standard errors and the dotted lines denote the read depth at which significance was tested. (c) Non-metric multidimensional scaling of bacterial compositions in samples based on Bray-Curtis dissimilarities. Samples are coloured by age and ellipses represent 95% confidence intervals. Arrows indicate the direction of cosines of the top 10 bacterial OTUs significantly correlated with the ordination axes, and are scaled by their strength of correlation (r2). (d) Mean DESeq2-transformed abundance of Actinobacteria and Firmicutes grouped by nutritional status and treatment. (e) Compositional variance among samples grouped by supplementation arm and age measured as distances to centroid, based on NMDS of weighted Unifrac dissimilarity scores. \**p* < 0.05, \*\**p* < 0.01, \*\*\**p* < 0.001 (f) Venn diagram showing the numbers of bacterial taxa with significantly increased or decreased abundance, as indicated by arrows, in supplemented groups relative to the control group. The pairs of numbers within brackets refer to taxa at 12 and 24 months of age respectively, and select taxa are listed in boxes. (g) Normalized abundance of *Escherichia-Shigella* and *Bifidobacterium* OTUs across supplementation arms at 12 months.

Analysis of compositional differences among samples revealed four distinct clusters of protozoan communities (Fig. 2e). The overall compositional variance was significantly explained by place of residence (adonis, R^2^ 0.02, *p* < 0.05) and micronutrient supplementation (adonis, R^2^ 0.09, *p* < 0.001), where protozoan communities in children supplemented with MNP differed significantly from those in control and MNP with zinc arms (MNP-CTL, R^2^ 0.05, *p* < 0.01; MNP-MNP with zinc R^2^ 0.04, *p* < 0.01). Cluster 1, in particular, was enriched in MNP samples, *X*^2^ (6, *N* = 114) = 38.5, *p* < 0.001 (Fig. 2f). Key drivers of the diversity included *Tritrichomonas*, detected almost exclusively in samples found in clusters 1 and 3 (correlation coefficient R^2^ 0.21, *p* = 0.001), and an OTU assigned to an unknown alveolate found predominantly in clusters 1 and 2 (R^2^ 0.17, *p* = 0.001). These organisms were highly prevalent in both age groups, at 42% and 45% (*Tritrichomonas*) and 20% and 21% (unknown alveolate). Fungal richness and phylogenetic composition were not associated with any of the variables studied here.

We identified significantly higher carriages of seven phylogenetically distinct protozoa and six fungi in children receiving MNPs without zinc, relative to those that were given zinc (six protozoa and six fungi relative to the control group; Fig. 2d). Indeed, we noted a trend where MNP with zinc reduced carriage of microbial eukaryotes to or below that observed in the control samples. For example, *Gregarina* and an uncharacterized alveolate, which contributed to the previously observed differences in beta diversity (Fig. 2e), were detected with 1.8 and 3.8-fold higher frequency in the MNP group, with no differences between samples from the control and MNP with zinc groups. Similarly, the carriages of three mucormycete genera (*Rhizomucor, Actinomucor* and *Mucor*) were 1.3, 1.5 and 1.8-fold higher, respectively, in the MNP group compared to the control, with no significant differences between the control and MNP with zinc groups. *Toxoplasma* was the only genus with significantly reduced carriage in children receiving MNP with zinc; however, we observed non-significant reductions in other organisms such as *Cercomonas* and *Mucor* (2 and 1.4-fold, respectively) suggesting possible species-specific effects. Despite previous reports of the impact of zinc on helminths^24^, we did not detect significant differences in the carriage of the tapeworm *Hymenolepis nana* among treatment arms.

### Micronutrient supplements are associated with specific bacterial communities

Using 16S rRNA amplicon sequencing, we also profiled the stool bacterial microbiota. From the 13,984,120 sequenced reads (median 92,628), we identified 1108 bacterial OTUs across all 160 samples (median 50; Supplementary Table 3). Actinobacteria and Firmicutes were found to dominate with just two OTUs (both assigned to *Bifidobacterium*) accounting for over 50% of all reads (Fig. 3a, Supplementary Table 4). Age was the primary determinant of bacterial richness (β = 43.65, CI [31.98 – 55.31], *p* < 0.001) and evenness (β = 0.80, CI [0.59 – 1.02], *p* < 0.001) (Fig. 3b, Supplementary Fig. 4,) as well as patterns of taxonomic composition as measured by Bray-Curtis and weighted Unifrac dissimilarities (Fig. 3c; adonis, R^2^ 0.06, *p* < 0.001; R^2^ 0.05, *p* < 0.001). Regression of dissimilarities in each child over time using partial correspondence analysis indicated that 56% of Bray-Curtis and 59% of weighted Unifrac changes may be attributed to age. By correlating the abundances of bacterial taxa with the first two axes of the Bray-Curtis ordination, we identified the candidate drivers of community differences as the two dominant *Bifidobacterium* species, with opposite abundance patterns perhaps suggesting succession of one species by the other.

**Fig. 4.**
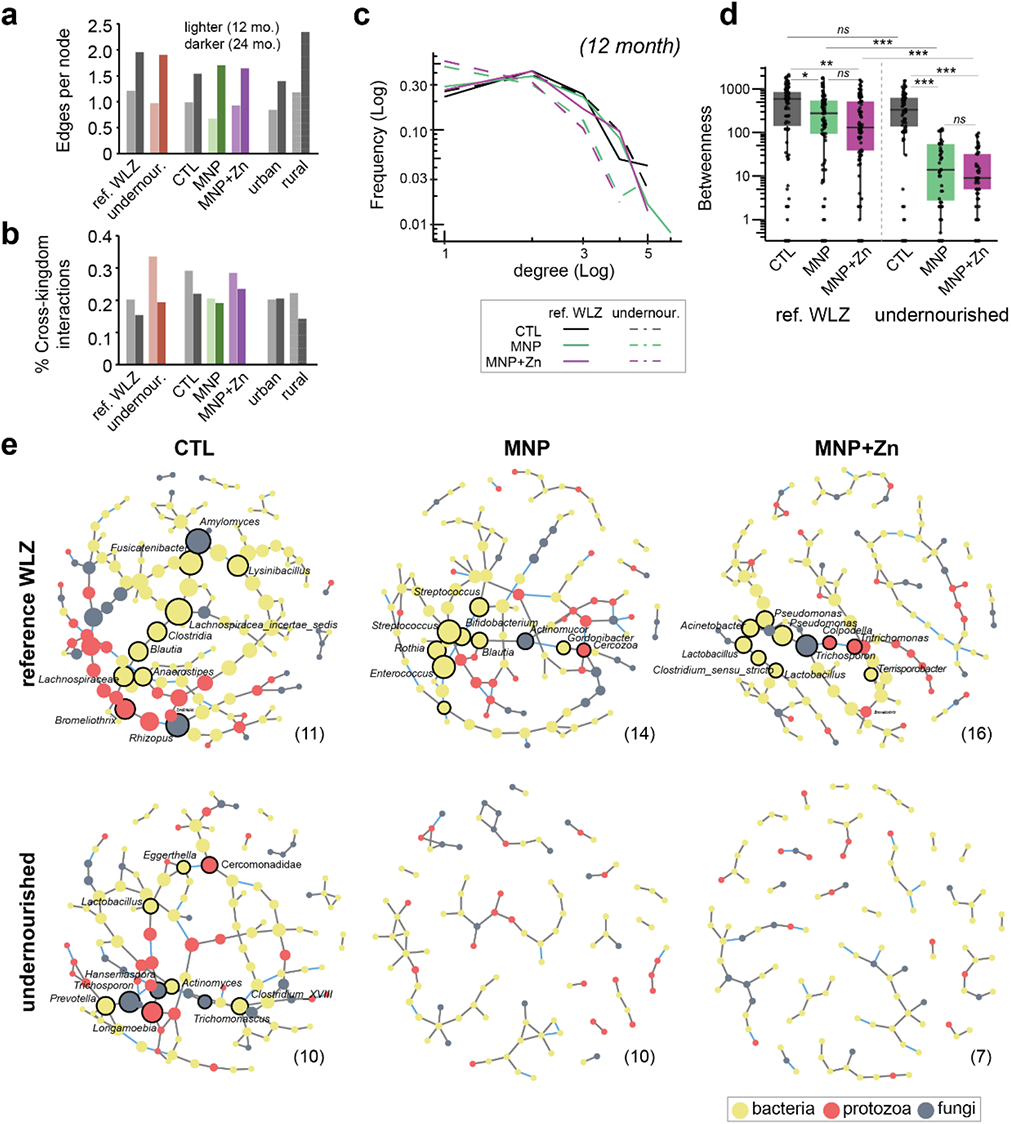
Supplementation influences microbial interactions. (a) Density of microbial interactions, calculated as significant correlations among microbiota (edges) normalized by the numbers of taxa (nodes), by nutritional status, supplementation arm and place of residence (site). Lighter and darker hues represent samples from 12 and 24 months respectively. (b) Proportions of significant microbial interactions occurring cross-kingdom, within indicated sample groups. (c) Degree distribution and (d) betweenness centrality scores of microbial networks in 12 month old children grouped by nutritional status and supplementation arm. (e) Graphic representations of aforementioned networks representing predicted microbial interactions in 12 month old children, grouped by nutritional status and micronutrient treatment. Nodes represent bacterial OTUs (yellow) and protozoan and fungal genera (red and grey, respectively), scaled by betweenness centrality scores. Edges represent significant positive (grey) and negative (blue) correlations among microbiota. Taxa with no predicted interactions have been removed. Numbers of samples used to generate each network are indicated within brackets.

Consistent with a previous study^27^, bacterial richness was reduced in undernourished children (β = −29.19, CI [−52.99 – −5.39], *p* < 0.05), while a significant interaction between nutritional status and place of residence indicated that bacterial evenness was reduced in undernourished children from the urban setting (β = 1.03, CI [0.11 – 1.95], *p* < 0.05) (Fig. 3b, Supplementary Fig. 4b). We detected no significant association between nutritional status and locality and bacterial beta diversities in this cohort.

Treatment with MNPs was associated with an overall increased abundance of Actinobacteria in children at 12 months compared to the control group and those receiving MNP with zinc (β = 36020, CI [7239 – 64802], *p* < 0.05), but reduced abundance in the MNP group at 24 months (β = −52670, CI [−93373 – −11966], *p* < 0.05) (Fig. 3d). Firmicutes were reduced in the presence of zinc in both age groups (β = −261976, CI [−476591 – −47362], *p* < 0.05), with a non-significant reduction in those supplemented without zinc (β = −206413, CI [−416049 – 3221], *p* = 0.055). Supplementation tended to reduce overall bacterial richness with an effect that reached significance in the MNP group (β = −14.66, CI [−29.01 – −0.31], *p* < 0.05) (Fig. 3b) and influenced taxonomic composition as measured by weighted Unifrac (adonis, R^2^ 0.03, *p* < 0.01) but not Bray-Curtis dissimilarities. Specifically, phylogenetic variance differed among groups (*p* < 0.001), with significantly smaller differences among 12 month old children receiving MNP and MNP with zinc (Tukey posthoc, *p* < 0.01) (Fig. 3e, Supplementary Fig. 4c). This may suggest that micronutrients support or restrict the growth of select taxa. Through differential abundance analysis, we identified 14 taxa with reduced abundances in both supplemented groups at 12 months compared to controls, including over 10-fold reductions in *Anaerostipes*, *Anaerosalibacter* and *Clostridium* XI (Fig. 3f). Two additional *Anaerostipes* OTUs were reduced in supplemented groups at both ages, with six OTUs reduced at 24 months only. MNP with zinc was associated with changes in an additional 46 taxa, and 29 taxa were altered in MNP samples. These included a seven-fold increase in *Escherichia-Shigella* abundance in 12 month old MNP-supplemented children, increases in several Lactobacilli and a 1.3-fold reduction in one *Bifidobacterium* OTU (Fig. 3g). These data reveal that micronutrient supplementation may impact bacterial communities during early development.

### MNPs may destabilize microbial interactions in undernourished infants

Microbial interaction networks were constructed to define significant taxonomic co-occurrences (Fig. 4). We found that interactions, calculated as edges per node, increased with age irrespective of treatment, nutritional status or place of residence, which reflects the development of more complex communities as the child matures (Fig. 4a). The greatest change, with a 2.5-fold increase, was noted in children in the MNP arm, which had the fewest taxon interactions at 12 months but achieved parity with the control and MNP with zinc groups by 24 months. Cross-kingdom interactions between bacteria and eukaryotes represented 20% to 30% of all interactions at 12 months, falling to between 15% and 24% by 24 months of age (Fig. 4b).

When split by nutritional status, we observed important differences in the networks of 12 month old undernourished infants supplemented with micronutrients compared to the control and reference WLZ groups (Fig. 4c,d). Within control groups, the microbial networks of undernourished infants and those within the reference WLZ group had similar levels of connectivity, with non-significant differences in degree distribution and betweenness centrality scores. While children in the reference WLZ group receiving either supplement were associated with small but significant reductions in microbiota betweenness (Wilcoxon rank sum, *p* < 0.05 and *p* < 0.01), greater reductions were observed in supplemented undernourished children (Wilcoxon rank sum, *p* < 0.001). Since betweenness provides a measure of the degree of coordination within a network, these findings suggest that micronutrient supplementation, with or without zinc, results in microbial communities that are less organized at 12 months of age. This is further illustrated by the network visualizations (Fig. 4e), where, in addition to changes in network density, we also identified shifts in taxa with the highest betweenness values (which can be interpreted as those taxa most likely to mediate important coordinating roles within the communities). For example, within the control group, Clostridia, two species of Mucoromycota and the ciliate *Bromeliothrix* occupy central roles in the network of reference WLZ infants, while in undernourished infants these central roles are held by *Trichosporon*, *Longamoeba* and *Prevotella*. In supplemented reference WLZ groups, Bacilli exhibit the highest betweenness values in the absence of zinc, while these are replaced by Proteobacteria in the communities from infants receiving MNP with zinc. However, within undernourished infants receiving either supplement, microbial networks appear largely fragmented (Fig. 4e), with dramatically lower degree distributions and betweenness compared to the control group suggesting that early treatment with micronutrient powders may destabilize a fragile microbial community. Comparison of microbial networks by location of residence further showed an increased density of interactions within each rural group (control or supplemented) compared to the urban groups (Supplementary Fig. 5). Low subject numbers precluded us from successfully generating networks at 24 months, where numbers of microbial taxa are greater.

**Fig. 5.**
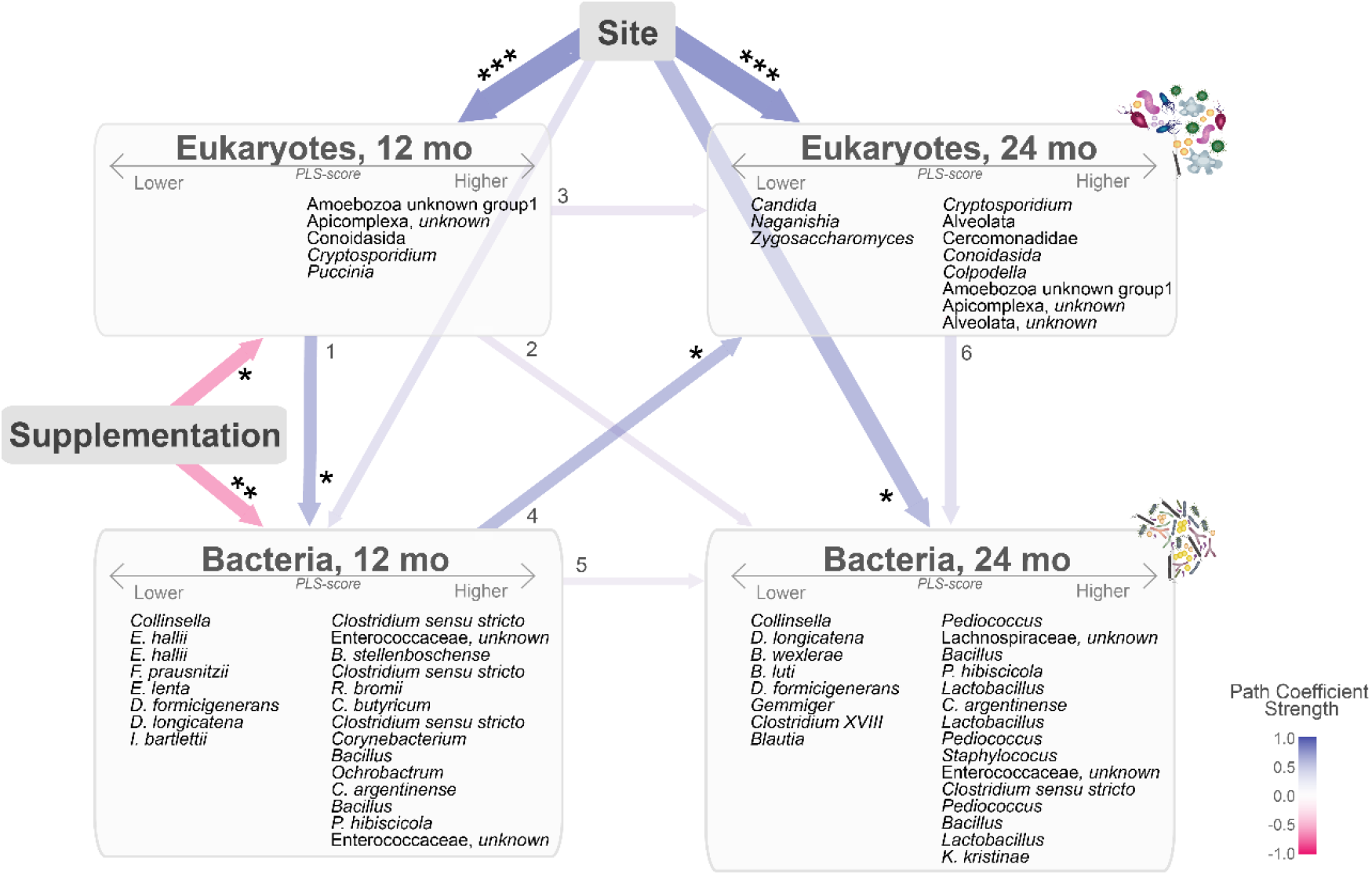
Graphic representation of the cross-associations among demographic variables, micronutrient supplementation and microbiota over time. Interconnected arrows indicate the tested cross-correlated paths between nodes of: place of residence (site), supplementation, and the composite measures of bacterial and eukaryotic OTUs detected at 12 and 24 months, collapsed as latent PLS-scores. Negative correlations are indicated in pink and positive in blue. Arrow thickness is weighted by the effect size of the direct path coefficients as indicated in Supplementary Table 5. Significance of direct paths, \**p* < 0.05, \*\**p* < 0.01, \*\*\**p* < 0.0001. OTUs that loaded positively (>0.4) or negatively (<−0.4) within each PLS-score are listed within boxes. PLS, partial least square; OTUs, operational taxonomic units.

### Complex cross-kingdom interrelationships over time are more influenced by place of residence than early supplementation

Based on our findings, we hypothesized that direct effects of supplementation and place of residence on microbial communities at 12 months could translate to indirect influence on later microbial profiles. We further hypothesized that early exposure to eukaryotes (before or at 12 months of age) would change the course of bacterial microbiome maturation. To explore the complex direct and indirect interrelationships among these factors, we generated an integrated model using partial least squares (PLS) path modelling (Fig. 5, Supplementary Table 5). First, place of residence had strong direct and indirect influences on eukaryotic and bacterial profiles at both 12 and 24 months. The greatest direct effects were on eukaryotic composition (12mo, path coefficient 0.52 ± 0.09, *p* < 0.0001; 24 mo, path coefficient 0.48± 0.1, *p* < 0.0001). Consistent with our findings above, children from the rural community had increased levels of several alveolates including *Cryptosporidium* at 12 and 24 months (12 months, 0.40 loading; 24 months 0.69 loading). While there was no significant direct effect on bacteria at 12 months (path coefficient 0.17 ± 0.12, *p* = 0.15), the locality indirectly influenced bacterial composition via eukaryotes (indirect path coefficient of 0.14 with a total effect of 0.31 at 12 months). Children from the rural community loaded positively for several *Clostridium* OTUs at both ages, and sustained higher levels of *Lactobacillus* at 24 months. Micronutrient supplementation appeared to influence the composition of eukaryotes and bacteria in an opposing manner to place of residence at 12 months (path coefficient −0.27 ± 0.11, *p* = 0.014; path coefficient −0.27 ± 0.09, *p* = 0.0058), with possible carryover effects to microbial compositions at 24 months (indirect effects of −0.11). Also consistent with our findings, *Mucor* and *Euglyphida* correlated with supplementation at 12 months (−0.35 and −0.34 cross-loadings, respectively).

Eukaryotic profiles at 12 months of age were significantly associated with a shift in bacterial profiles at 12 months suggesting possible cross-kingdom interactions (Fig. 5, arrow 1; path coefficient 0.27 ± 0.12, *p* = 0.033). These bacteria, in turn, exhibited a significant influence on eukaryotic composition at 24 months (Fig. 5, arrow 4; path coefficient 0.21 ± 0.095, *p* = 0.033). Differences in path coefficients were also tested in a stratified analysis of reference WLZ and undernourished children but none reached statistical significance in our cohort. While the pathway coefficients identified above were found to be statistically significant, due to large standard errors likely resulting from heterogeneity and small sample size, we were unable to validate this support using more robust bootstrapping procedures (Supplementary Table 5). Nevertheless, given the consistency of these relationships with our earlier findings, this model provides additional support for the indirect association of MNP supplementation and bacterial communities mediated through the promotion of specific eukaryotic microbes.

## Discussion

Malnutrition, both undernutrition and obesity, are associated with altered bacterial compositions, where in the former, underdeveloped bacterial communities have the capacity to induce weight loss^6,28^. Here, we have shown that the gut microbiota of both undernourished children and those within a healthy weight range include a diverse group of protozoa, helminths and fungi, each with the capacity to impact host health. We have also shown that supplementation with MNPs, a strategy used to improve growth and alleviate micronutrient deficiencies^14,16^, has the capacity to influence the development of the microbiome in these susceptible populations.

Consistent with previous studies, we found that bacterial communities became more complex during growth. Eukaryotic communities, however, were not significantly impacted by age, but instead were associated with micronutrient supplementation and place of residence. Only the tapeworm *H. nana* was identified at significantly higher levels in undernourished children. While *H. nana* infection is usually asymptomatic, high egg burdens in children have previously been associated with diarrhea, abdominal pain and weight loss^29^, with exacerbated morbidity in children <5 years^30^. We associated rural habitation with significantly more diverse protozoan communities, and in particular increased prevalence of *Cryptosporidium*. An important cause of infant mortality and childhood malnutrition, *Cryptosporidium* infection is attributed to unsafe drinking water and inadequate sanitation often associated with rural settings^26,31^. While approximately half of all children enrolled in the trial had access to piped drinking water (41% and 52% in the urban Bilal colony and rural Matiari sites respectively), only 4% of children in the Matiari district had access to underground sewage, compared to 95% in the Bilal Colony^26^, consistent with a lack of waste water sanitation resulting in higher parasite carriage. While the large multicenter GEMS study reported *Cryptosporidium* as a leading cause of death in 12 to 23 month old children with moderate to severe diarrhea in developing countries^32^, we found a high prevalence of this parasite in absence of diarrhea (80% and 83% at 12 and 24 months in the Matiari district, and 60% and 33% in the Bilal urban colony). As our detection is based on 18S rRNA amplicon sequencing, we may have detected a broader group of species of variable pathogenic potential compared to the GEMS study, which applied a specific oocyst antigen immunoassay. Alternatively, our findings may indicate a high prevalence of asymptomatic infections, with symptomatic infections resulting from additional unknown factors^7,33^. The prevalence of *Cryptosporidium* in our cohort was also higher than previously reported in non-diarrheal stools, using oocyst antigen testing, in the neighbouring Naushero Feroze District (5.1% between 12 and 21 months of age), where *Cryptosporidium* contributed to 8.8 diarrheal episodes per 100 child years^34,35^. This same study associated asymptomatic enteropathogen infection, including *Cryptosporidium* and *Giardia*, across eight countries with elevated inflammation and intestinal permeability, factors thought to increase risk of stunting and impact the effectiveness of nutritional interventions in low-resource settings^35^.

A major focus of our study was to estimate the effect of micronutrient supplementation on the gut microbiota. We found that children receiving supplements without zinc were associated with distinct eukaryotic communities, featuring an increased prevalence of multiple protozoan and fungal taxa; however, the addition of zinc to these supplements alleviated these increases, while significantly reducing the prevalence of *Toxoplasma* and overall protozoan richness. These findings are consistent with a previous report which suggested that zinc has a parasite-specific protective effect against infection and ensuing diarrhea^24^. Fungal diversity was not impacted by age, supplementation, place of residence or nutritional status. However, the predominance of Mucoromycota, particularly in children receiving MNPs without zinc, is of concern, as these organisms are responsible for rare but lethal invasive fungal infections that have previously been reported in low birth weight infants and malnourished children^36^. Although incidence of infections is rising globally, rates of mucormycoses are particularly high in Asia^37^. Notably, a recent spike in infections, also termed ‘Black fungus’, in thousands of active and recovered Covid-19 patients in India, was attributed to treatment with corticosteroids to control inflammation, in conjunction with a high prevalence of diabetes^38^.

It has been well established that iron supplementation can promote the virulence of particular fungi and parasites^39,40^. Several studies have shown that iron alone or in combination with other micronutrients worsens existing infections, lengthens the duration and severity of diarrhea and increases mortality rates in children^22,26,39^. Consequently, sequestration of free iron by host proteins such as lactoferrin is a key defense mechanism to limit growth of pathogens including Mucorales^41^. Iron deficiency has furthermore been suggested as protective against malaria infection^42,43^, and provision of supplements containing iron in endemic regions has been cautioned against due to increased malaria-related hospitalization and mortality of children^39^. While deficiency in zinc has been associated with impaired immune function and susceptibility to enteroinfections^44^, supplementation in the context of enteric pathogens was shown to have parasite-specific outcomes. Provision of zinc alone can increase the incidence of *Ascaris lumbricoides* and duration of *Entamoeba histolytica* infections, but it has also been shown to reduce the duration of associated diarrheal episodes as well as lower the prevalence of *Giardia lamblia* infections^24^. Interestingly, asymptomatic *Giardia* infections in children in Tanzania were associated with reduced rates of diarrhea and fever, an effect which was lost in children receiving vitamin and mineral supplements, including both iron and zinc^45^. Our data suggest that while iron, vitamins, or both, may promote growth and survival of commensal and potentially pathogenic eukaryotes, resulting in a shift in eukaryotic community structure, the addition of zinc may reduce the ability of at least some eukaryotic microbes to infect and persist. The findings of reduced bacterial diversity in 12 month old infants receiving micronutrient supplements, together with elevated levels of *Escherichia-Shigella* and reduced beneficial *Bifidobacteria*, are also consistent with previous reports, where reductions in beneficial *Bifidobacterium* and *Lactobacilli* and increased enterobacteria in infants receiving iron-containing micronutrients were linked to elevated risk of inflammation and diarrhea^22,23,46^. The original cRCT trial associated *Aeromonas* infection with increased diarrhea in MNP supplemented groups^26^. We did not detect this bacterium in our data, possibly due to exclusion of diarrheal samples.

The impact of micronutrient supplementation also extended to the structure of the microbial communities. Microbial networks, representing significant correlations in the co-occurrence of bacteria and eukaryotes, revealed higher network connectivity in the control groups, with the networks generated from the undernourished infants receiving both types of supplements, revealing a more fragmented structure. This fragmentation suggests a destabilization of species-interactions within the developing gut microbiota in undernourished infants. Possibly contributing to this destabilization is the presence of specific eukaryotic microbes, as evidenced by higher proportions of eukaryotic-bacterial interactions in healthy infants receiving either supplement, and/or the expansion of pathogenic bacteria. These microbes may interfere with the maturation of commensal bacteria through predation, competition for resources and/or modulation of host immunity. In undernourished infants, the cumulative effect of increases in pathogenic organisms on community structure may be more pronounced than in infants within a healthy weight. Enteropathogens *Giardia lamblia* and enteroaggregative *Escherichia coli*, for example, were shown to have a greater impact on growth in protein-deficient mice during co-infection, an effect which was dependent on the resident gut bacteria^47^. Taken together, our data showing increased carriage of eukaryotic microbes and increased abundance of *Escherichia-Shigella* in children supplemented with micronutrients, as well as a potential loss of organization in microbial interactions in supplemented undernourished children, may offer at least a partial explanation for previous reports of increased duration and severity of diarrhea as well as increased intestinal inflammation in children supplemented with micronutrient powders^26^.

Due to the relatively small numbers of samples, we were unable to generate separate networks for the three treatment arms for 24 month old children. We note that supplementation had ceased six months prior, consequently the acute effects of these supplements may have dissipated. Small sample sizes also preclude us from further segregating microbial networks by place of residence. Micronutrient interventions may impact undernourished children differently in the context of a high *Cryptosporidium* burden, for example. The notable absence of *Giardia*, a parasite typically prevalent in this demographic, is likely due to mismatches to the 18S rRNA sequencing primers^13^. Nevertheless, parasite diagnostic data from the trial did identify *Giardia* in 37 infants at 12 months, and *Cryptosporidium* in seven, but noted no significant increases in either of the supplemented groups^26^. Prevalence was nearly two-fold higher at the rural site, consistent with our findings for *Cryptosporidium*, emphasizing the need for location-specific investigations of the effects of micronutrient supplements. In addition to potential intraspecies variation, our detection of high sequence diversity in *Cryptosporidium* OTUs specifically, and eukaryotic taxa in general, may be exaggerated by a high proportion of non-overlapping amplicon reads, a consequence we have attempted to minimize through manual curation. Regardless, we report that eukaryotic microbiota are abundant members of the gut microbiome even in infancy, and given the known role of parasitic pathogens in diarrheal disease and the association of fungi with obesity and inflammatory bowel disease^48,49^, their role in malnutrition should be further studied.

Although not supported by robust bootstrapping, our integrated model of microbial relationships and influencing external factors was able to recapitulate a number of key earlier findings, including the impact of locality and micronutrients on gut eukaryotes. Furthermore, the prediction from our model that complex cross-kingdom interactions may influence gut bacterial composition, provides a valuable framework to dissect the direct and indirect effects of eukaryotic infections or nutritional interventions on the maturing gut microbiome. Given the current debate over the use of MNP supplementation and its role in gastrointestinal disorders, such a framework is expected to play a key role in identifying scenarios where MNP supplementation may require more cautious thinking.

## Conclusion

This study demonstrates that micronutrient powders impact the infant microbiota, with potentially destabilizing effects driven through the promotion of specific organisms during early stages of microbiome development. These findings are of relevance to micronutrient supplementation strategies, especially those targeting vulnerable children in low resource settings.

## Methods

### Study design and subject selection

Study participants were selected from a multicenter clustered randomized controlled trial (ClinicalTrials.gov identifier NCT00705445) that investigated the effects of micronutrient supplementation with or without zinc among 2746 children from either an urban (Bilal colony, squatter settlement within Karachi) or rural (Matiari district, 200 km from Karachi) site in Sindh, Pakistan^26^. In the trial, daily supplementation with micronutrient powders (MNP) containing vitamins A, C, D, folic acid and microencapsulated iron, with or without zinc spanned 6 to 18 months of age, with prospective follow-up until 24 months for the collection of health and demographic information and stool samples^26^. Eighty children were selected for microbiome profiling according to the following criteria (Supplementary Fig. 1): 1) having stool samples collected at 12 and 24 months of age available and archived at −80°C; 2) having at 24 months a weight-for-length z-score (WLZ) < −2 below the median (undernourished) or > −1 (reference WLZ) based on WHO 2006 growth references (www.who.int/childgrowth); 3) no record of antibiotic administration within 14 days of stool sample collection; and, 4) no reported diarrhea within seven days of stool collection. Subjects within the reference group were further selected based on fewest WLZ scores < −1 at other time points, to represent as healthy as possible a comparator group. Participant characteristics were summarized as medians with interquartile ranges (IQRs) or means ± standard deviations (SD) if continuous variables, and percentages if categorical.

### DNA extraction and amplicon sequencing

DNA was extracted from 100-200 mg of stool using the E.Z.N.ATM Stool kit (Omega Bio-Tek Inc, GA, USA) according to the manufacturer’s protocol. Mechanical disruption of cells was carried out with the MP Bio FastPrep-24 for 5 cycles of 1 min at 5.5 M/s. 16S variable region 4 (V4) amplifications were carried out using the KAPA2G Robust HotStart ReadyMix (KAPA Biosystems) and barcoded primers 515F and 806R^50^. The cycling conditions were 95°C for 3 min, 22 cycles of 95°C for 15 s, 50°C for 15 s and 72°C for 15 s, followed by a 5 min 72°C extension. Libraries were purified using Ampure XP beads and sequenced using MiSeq V2 (150bp x 2) chemistry (Illumina, San Diego, CA). 18S V4+V5 amplification was achieved using the iProof DNA polymerase (Bio-Rad Laboratories, Hercules, CA) with primers V4-1 and V4-4 as previously described^13^. Briefly, the cycling conditions used were 94°C for 3 min, 30 cycles of 94°C for 45 s, 56°C for 1 min and 72°C for 1 min, followed by a 10 min 72°C extension. Barcodes were ligated and libraries were sequenced using MiSeq V3 (300bp x 2) chemistry (Illumina, San Diego, CA). Sequencing was performed at the Centre for the Analysis of Genome Evolution and Function (Toronto, Canada).

### Sequence data analysis

16S data were quality filtered and processed using VSEARCH v2.10.4^51^ and the UNOISE pipeline in USEARCH v11.0.667^52,53^. Filtered sequences were clustered to 99% sequence identity, and the resulting operational taxonomic units (OTUs) were classified with a minimum confidence of 0.8 using the SINTAX^54^ algorithm and the Ribosomal Database Project version 16^55^.

18S data were quality filtered using Trimmomatic v0.36^56^ and read pairs with minimum 200 nucleotide length were merged using VSEARCH, or artificially joined using a linker of 50 ambiguous nucleotides (N50) using USEARCH. Resultant amplicon sequences were clustered to 97% sequence identity using the UCLUST^52^ algorithm, and taxonomically classified using SINA v1.2.11^57^ with a minimum 90% sequence similarity threshold. Unclassified sequences were submitted for classification using SINTAX and the SILVA v132 non-redundant reference database^58^, and those still unclassified were compared to the NCBI non-redundant nucleotide database^59^ (downloaded Nov 28, 2017) by BLAST^60^ using a 90% cutoff for both sequence identity and query coverage. Phylogenetic tree construction for both 16S and 18S OTUs was performed using the FastTree^61^ algorithm and visualized using the Iroki viewer^62^, with taxon prevalence values calculated at a minimum threshold of 5 reads.

### Microbial diversity and differential abundance analyses

Microbiota richness (number of OTUs) and evenness (Shannon Diversity Index, H) were calculated using Phyloseq 1.20.0^63^. Rarefaction curves were generated at 100 read intervals to a maximum of 5,000 or 50,000 for eukaryotes and bacteria, respectively. Values were averaged and standard errors calculated by the grouping variable. As intra-class correlation was low, we implemented generalized linear models (GLMs) using richness and evenness values averaged from 100 independent rarefactions at read depths of 25,000 (bacteria) and 1,000 (protozoa and fungi). To identify a final model that best explains diversity, we performed stepwise model selection using AIC with MASS^64^ with the following explanatory variables: age, nutritional status, supplementation and urban versus rural site.

Differences in bacterial composition, based on Bray-Curtis and weighted Unifrac dissimilarity scores, were calculated with Phyloseq and vegan^65^ using DESeq2-normalized counts prefiltered for taxa represented by a minimum of 5 reads in at least 5% of the samples. The contribution of age to beta diversity was calculated using the capscale function, and the remaining variables were tested for significance in age-stratified samples using adonis. The compositional variance within groups, measured as distances to centroids, was evaluated using the betadisper function, and pairwise differences were delineated using a post hoc Tukey test. All adonis and betadisper tests were carried out with 9999 permutations. We applied non-metric dimensional scaling (NMDS) to ordinate samples based on their compositional dissimilarity. The envfit function was used to identify taxa significantly correlated with the first two ordination axes (candidate drivers of community differences), indicated by arrows in the direction of cosines and scaled by the root square of the correlation. Protozoan and fungal beta diversities were evaluated at 1000 read depth using Principal Coordinate Analysis of unweighted Unifrac scores, and significance was tested as above. Differential taxon abundance was tested with DESeq2 1.22.2^66^ in samples containing a minimum of 1000 reads, using data internally transformed with the median of ratios method.

Fisher’s Exact or pairwise test from the rstatix package was used to evaluate differences in eukaryote carriage among participant groups, using a minimum 5 read detection threshold per OTU and grouping OTUs to the genus level or the lowest assigned taxonomic level. Benjamini-Hochberg correction was applied for multiple testing.

### Microbial interaction networks

Bacterial and eukaryotic datasets were rarefied to 25,000 and 1,000 reads, respectively, and eukaryotes were agglomerated to genera or the lowest assigned taxonomic level. Microbial interaction networks, including both microbial datasets simultaneously, were generated using SpiecEasi^67^ with the neighbour selection (MB) method, nlambda 100 and lambda.min.ratio 1e-02, and visualized using igraph^68^.

### Partial least squares path analysis

To explore the complex system of direct and indirect relationships between micronutrient supplementation, place of residence and the multivariate matrices of bacteria and eukaryotes over time, we conducted partial least squares (PLS) path analysis using the plspm package in R^69^. Microbial read counts were center-log transformed after pre-filtering for taxa with more than 0.01% abundance across all samples. The analysis was set to collapse the high dimensional microbial community matrices into latent PLS-scores representing community patterns of 1) eukaryotes at 12 months, 2) eukaryotes at 24 months, 3) bacteria at 12 months and 4) bacteria at 24 months. The analysis estimates the relationships between factors based on cross correlations, e.g. how eukaryotes detected at 12 months load into a community pattern summarized by a latent PLS-score (i.e. “Eukaryotes, 12 mo”) in a manner that optimises the cross-correlation with the other variables (i.e. supplementation, place of residence and other community patterns). Path coefficients indicate the strength of the internodal relationship and can be conceptually understood as correlation coefficients. Bootstrapping procedures were followed for validation and differences in path coefficients were also tested between nutritional groups.

All microbial data and statistical analyses were carried out with R version 4.0.2^70^.

### Ethics Approval

The protocol for the cRCT trial was approved by the Ethics Review Committee of Aga Khan University (752-Peds/ERC-07). This sub-study protocol was approved by research ethics board at The Hospital for Sick Children, Toronto (REB No. 1000054244), the ethics review committee at Aga Khan University, Karachi, Pakistan (4840-Ped-ERC-17), and the National Bioethics Committee Pakistan (4-87/NBC-277/17/1191).

## Data availability

Raw sequence data have been deposited to the NCBI Sequence Read Archive with the BioProject identifier PRJNA717317.

## Code availability

R code for analyses is available on GitHub (https://github.com/ParkinsonLab/gut-eukaryotes-malnutrition-and-micronutrient-supplementation).

## Acknowledgements

We thank Imran Ahmed (Aga Khan University, Karachi, Pakistan) and Didar Alam (Aga Khan University) for assistance with organizing stool sample shipments from Pakistan to Canada and providing secure access to clinical data. We also appreciate the helpful advice and insights from Amel Taibi and Elena Comelli in addressing challenges encountered during extraction of sample DNA. This work was supported by a HSBC Bank Canada Catalyst Research Grant from the Hospital for Sick Children awarded to CB, RB, DG, JP and LGP; the Canadian Institute for Health Research grant PJT-152921 to JP; Restracomp scholarship administered by the Research Training Centre (Hospital for Sick Children) and a graduate scholarship from the Government of Ontario to AP. Computing resources were provided by the SciNet High Performance Computing (HPC) Consortium; SciNet is funded by the Canada Foundation for Innovation under the auspices of Compute Canada, the Government of Ontario, Ontario Research Fund - Research Excellence, and the University of Toronto.

## Author contributions

L.G.P., Z.A.B., J.P. and R.H.J.B. conceived and designed the study. S.S. and Z.A.B. participated in original collection of clinical samples. A.P. isolated DNA and processed the sequencing data.

P.W.W and D.S.G. aided in design of amplicon generation. A.P. and C.B. analyzed the data and wrote the paper and all authors reviewed and/or edited the paper.

## References

1 Global Nutrition Report: Action on equity to end malnutrition., (Bristol, UK, 2020).

2 (UNICEF), U. N. C. s. F., Organization, W. H. & Bank, I. B. f. R. a. D. T. W. Levels and trends in child malnutrition: key findings of the 2021 edition of the joint child malnutrition estimates., (Geneva, 2021).

3 Black, R. E. et al. Maternal and child undernutrition and overweight in low-income and middle-income countries. Lancet 382, 427–451, doi:10.1016/S0140-6736(13)60937-X (2013).

4 Liu, L. et al. Global, regional, and national causes of under-5 mortality in 2000-15: an updated systematic analysis with implications for the Sustainable Development Goals. Lancet 388, 3027–3035, doi:10.1016/S0140-6736(16)31593-8 (2016).

5 Subramanian, S. et al. Persistent gut microbiota immaturity in malnourished Bangladeshi children. Nature 510, 417–421, doi:10.1038/nature13421 (2014).

6 Smith, M. I. et al. Gut microbiomes of Malawian twin pairs discordant for kwashiorkor. Science 339, 548–554, doi:10.1126/science.1229000 (2013).

7 Desai, N. T., Sarkar, R. & Kang, G. Cryptosporidiosis: An under-recognized public health problem. Trop Parasitol 2, 91–98, doi:10.4103/2229-5070.105173 (2012).

8 Mondal, D. et al. Contribution of enteric infection, altered intestinal barrier function, and maternal malnutrition to infant malnutrition in Bangladesh. Clin Infect Dis 54, 185–192, doi:10.1093/cid/cir807 (2012).

9 Ramanan, D. et al. Helminth infection promotes colonization resistance via type 2 immunity. Science 352, 608–612, doi:10.1126/science.aaf3229 (2016).

10 Chappell, C. L. et al. Fecal Indole as a Biomarker of Susceptibility to Cryptosporidium Infection. Infect Immun 84, 2299–2306, doi:10.1128/IAI.00336-16 (2016).

11 Chudnovskiy, A. et al. Host-Protozoan Interactions Protect from Mucosal Infections through Activation of the Inflammasome. Cell 167, 444–456 e414, doi:10.1016/j.cell.2016.08.076 (2016).

12 Reynolds, L. A. et al. Commensal-pathogen interactions in the intestinal tract: lactobacilli promote infection with, and are promoted by, helminth parasites. Gut Microbes 5, 522–532, doi:10.4161/gmic.32155 (2014).

13 Popovic, A. et al. Design and application of a novel two-amplicon approach for defining eukaryotic microbiota. Microbiome 6, 228, doi:10.1186/s40168-018-0612-3 (2018).

14 Tam, E., Keats, E. C., Rind, F., Das, J. K. & Bhutta, A. Z. A. Micronutrient Supplementation and Fortification Interventions on Health and Development Outcomes among Children Under-Five in Low- and Middle-Income Countries: A Systematic Review and Meta-Analysis. Nutrients 12, doi:10.3390/nu12020289 (2020).

15 Keats, E. C. et al. Effective interventions to address maternal and child malnutrition: an update of the evidence. Lancet Child Adolesc Health 5, 367–384, doi:10.1016/S2352-4642(20)30274-1 (2021).

16 (UNICEF), U. N. C. s. F. Nutrition, for Every Child: UNICEF Nutrition Strategy 2020–2030., (UNICEF, New York, 2020).

17 Mayo-Wilson, E. et al. Zinc supplementation for preventing mortality, morbidity, and growth failure in children aged 6 months to 12 years of age. Cochrane Database Syst Rev, CD009384, doi:10.1002/14651858.CD009384.pub2 (2014).

18 Imdad, A., Mayo-Wilson, E., Herzer, K. & Bhutta, Z. A. Vitamin A supplementation for preventing morbidity and mortality in children from six months to five years of age. Cochrane Database Syst Rev 3, CD008524, doi:10.1002/14651858.CD008524.pub3 (2017).

19 Papier, K. et al. Childhood malnutrition and parasitic helminth interactions. Clin Infect Dis 59, 234–243, doi:10.1093/cid/ciu211 (2014).

20 Hibberd, M. C. et al. The effects of micronutrient deficiencies on bacterial species from the human gut microbiota. Sci Transl Med 9, doi:10.1126/scitranslmed.aal4069 (2017).

21 Becker, K. W. & Skaar, E. P. Metal limitation and toxicity at the interface between host and pathogen. FEMS Microbiol Rev 38, 1235–1249, doi:10.1111/1574-6976.12087 10.1111/1574-6976.12087. Epub 2014 Sep 29. (2014).

22 Jaeggi, T. et al. Iron fortification adversely affects the gut microbiome, increases pathogen abundance and induces intestinal inflammation in Kenyan infants. Gut 64, 731–742, doi:10.1136/gutjnl-2014-307720 (2015).

23 Paganini, D. et al. Iron-containing micronutrient powders modify the effect of oral antibiotics on the infant gut microbiome and increase post-antibiotic diarrhoea risk: a controlled study in Kenya. Gut 68, 645–653, doi:10.1136/gutjnl-2018-317399 (2019).

24 Long, K. Z. et al. Effect of vitamin A and zinc supplementation on gastrointestinal parasitic infections among Mexican children. Pediatrics 120, e846–855, doi:10.1542/peds.2006-2187 (2007).

25 Richard, S. A. et al. Zinc and iron supplementation and malaria, diarrhea, and respiratory infections in children in the Peruvian Amazon. Am J Trop Med Hyg 75, 126–132, doi:10.4269/ajtmh.2006.75.1.0750126 (2006).

26 Soofi, S. et al. Effect of provision of daily zinc and iron with several micronutrients on growth and morbidity among young children in Pakistan: a cluster-randomised trial. The Lancet 382, 29–40, doi:10.1016/S0140-6736(13)60437-7 (2013).

27 Mendez-Salazar, E. O., Ortiz-Lopez, M. G., Granados-Silvestre, M. L. A., Palacios-Gonzalez, B. & Menjivar, M. Altered Gut Microbiota and Compositional Changes in Firmicutes and Proteobacteria in Mexican Undernourished and Obese Children. Front Microbiol 9, 2494, doi:10.3389/fmicb.2018.02494 (2018).

28 Ridaura, V. K. et al. Gut microbiota from twins discordant for obesity modulate metabolism in mice. Science 341, 1241214, doi:10.1126/science.1241214 (2013).

29 Ryan, E. T., Hill, D. R., Solomon, T., Endy, T. P. & Aronson, N. Hunter’s tropical medicine and emerging infectious diseases. (2020).

30 Soares Magalhaes, R. J. et al. Extending helminth control beyond STH and schistosomiasis: the case of human hymenolepiasis. PLoS Negl Trop Dis 7, e2321, doi:10.1371/journal.pntd.0002321 (2013).

31 Daniels, M. E., Smith, W. A. & Jenkins, M. W. Estimating Cryptosporidium and Giardia disease burdens for children drinking untreated groundwater in a rural population in India. PLoS Negl Trop Dis 12, e0006231, doi:10.1371/journal.pntd.0006231 (2018).

32 Kotloff, K. L. et al. Burden and aetiology of diarrhoeal disease in infants and young children in developing countries (the Global Enteric Multicenter Study, GEMS): a prospective, case-control study. Lancet 382, 209–222, doi:10.1016/S0140-6736(13)60844-2 (2013).

33 Ajjampur, S. S. et al. Symptomatic and asymptomatic Cryptosporidium infections in children in a semi-urban slum community in southern India. Am J Trop Med Hyg 83, 1110–1115, doi:10.4269/ajtmh.2010.09-0644 (2010).

34 Platts-Mills, J. A. et al. Use of quantitative molecular diagnostic methods to assess the aetiology, burden, and clinical characteristics of diarrhoea in children in low-resource settings: a reanalysis of the MAL-ED cohort study. Lancet Glob Health 6, e1309–e1318, doi:10.1016/S2214-109X(18)30349-8 (2018).

35 Kosek, M. N. & Investigators, M.-E. N. Causal Pathways from Enteropathogens to Environmental Enteropathy: Findings from the MAL-ED Birth Cohort Study. EBioMedicine 18, 109–117, doi:10.1016/j.ebiom.2017.02.024 (2017).

36 Francis, J. R., Villanueva, P., Bryant, P. & Blyth, C. C. Mucormycosis in Children: Review and Recommendations for Management. J Pediatric Infect Dis Soc 7, 159–164, doi:10.1093/jpids/pix107 (2018).

37 Prakash, H. & Chakrabarti, A. Global Epidemiology of Mucormycosis. J Fungi (Basel) 5, doi:10.3390/jof5010026 (2019).

38 Raut, A. & Huy, N. T. Rising incidence of mucormycosis in patients with COVID-19: another challenge for India amidst the second wave? Lancet Respir Med, doi:10.1016/S2213-2600(21)00265-4 (2021).

39 Sazawal, S. et al. Effects of routine prophylactic supplementation with iron and folic acid on admission to hospital and mortality in preschool children in a high malaria transmission setting: community-based, randomised, placebo-controlled trial. Lancet 367, 133–143, doi:10.1016/S0140-6736(06)67962-2 (2006).

40 Symeonidis, A. S. The role of iron and iron chelators in zygomycosis. Clin Microbiol Infect 15 Suppl 5, 26–32, doi:10.1111/j.1469-0691.2009.02976.x (2009).

41 Ibrahim, A. S., Spellberg, B., Walsh, T. J. & Kontoyiannis, D. P. Pathogenesis of mucormycosis. Clin Infect Dis 54 Suppl 1, S16–22, doi:10.1093/cid/cir865 (2012).

42 Gwamaka, M. et al. Iron deficiency protects against severe Plasmodium falciparum malaria and death in young children. Clin Infect Dis 54, 1137–1144, doi:10.1093/cid/cis010 (2012).

43 Jonker, F. A. et al. Iron status predicts malaria risk in Malawian preschool children. PLoS One 7, e42670, doi:10.1371/journal.pone.0042670 (2012).

44 Shankar, A. H. & Prasad, A. S. Zinc and immune function: the biological basis of altered resistance to infection. Am J Clin Nutr 68, 447S–463S, doi:10.1093/ajcn/68.2.447S (1998).

45 Veenemans, J. et al. Protection against diarrhea associated with Giardia intestinalis Is lost with multi-nutrient supplementation: a study in Tanzanian children. PLoS Negl Trop Dis 5, e1158, doi:10.1371/journal.pntd.0001158 (2011).

46 Paganini, D. & Zimmermann, M. B. The effects of iron fortification and supplementation on the gut microbiome and diarrhea in infants and children: a review. Am J Clin Nutr 106, 1688S–1693S, doi:10.3945/ajcn.117.156067 (2017).

47 Bartelt, L. A. et al. Cross-modulation of pathogen-specific pathways enhances malnutrition during enteric co-infection with Giardia lamblia and enteroaggregative Escherichia coli. PLoS Pathog 13, e1006471, doi:10.1371/journal.ppat.1006471 (2017).

48 Sokol, H. et al. Fungal microbiota dysbiosis in IBD. Gut 66, 1039–1048, doi:10.1136/gutjnl-2015-310746 (2017).

49 Mar Rodriguez, M. et al. Obesity changes the human gut mycobiome. Sci Rep 5, 14600, doi:10.1038/srep14600 (2015).

50 Caporaso, J. G. et al. Global patterns of 16S rRNA diversity at a depth of millions of sequences per sample. Proc Natl Acad Sci U S A 108 Suppl 1, 4516–4522, doi:10.1073/pnas.1000080107 (2011).

51 Rognes, T., Flouri, T., Nichols, B., Quince, C. & Mahe, F. VSEARCH: a versatile open source tool for metagenomics. PeerJ 4, e2584, doi:10.7717/peerj.2584 (2016).

52 Edgar, R. C. Search and clustering orders of magnitude faster than BLAST. Bioinformatics 26, 2460–2461, doi:10.1093/bioinformatics/btq461 (2010).

53 Edgar, R. C. UNOISE2: improved error-correction for Illumina 16S and ITS amplicon sequencing. bioRxiv (2016).

54 Edgar, R. C. SINTAX: a simple non-Bayesian taxonomy classifier for 16S and ITS sequences. bioRxiv (2016).

55 Cole, J. R. et al. Ribosomal Database Project: data and tools for high throughput rRNA analysis. Nucleic Acids Res 42, D633–642, doi:10.1093/nar/gkt1244 (2014).

56 Bolger, A. M., Lohse, M. & Usadel, B. Trimmomatic: a flexible trimmer for Illumina sequence data. Bioinformatics 30, 2114–2120, doi:10.1093/bioinformatics/btu170 (2014).

57 Pruesse, E., Peplies, J. & Glockner, F. O. SINA: accurate high-throughput multiple sequence alignment of ribosomal RNA genes. Bioinformatics 28, 1823–1829, doi:10.1093/bioinformatics/bts252 (2012).

58 Quast, C. et al. The SILVA ribosomal RNA gene database project: improved data processing and web-based tools. Nucleic Acids Res 41, D590–596, doi:10.1093/nar/gks1219 (2013).

59 Coordinators, N. R. Database Resources of the National Center for Biotechnology Information. Nucleic Acids Res 45, D12–D17, doi:10.1093/nar/gkw1071 (2017).

60 Altschul, S. F., Gish, W., Miller, W., Myers, E. W. & Lipman, D. J. Basic local alignment search tool. J Mol Biol 215, 403–410, doi:10.1016/S0022-2836(05)80360-2 (1990).

61 Price, M. N., Dehal, P. S. & Arkin, A. P. FastTree: computing large minimum evolution trees with profiles instead of a distance matrix. Mol Biol Evol 26, 1641–1650, doi:10.1093/molbev/msp077 (2009).

62 Moore, R. M., Harrison, A. O., McAllister, S. M., Polson, S. W. & Wommack, K. E. Iroki: automatic customization and visualization of phylogenetic trees. PeerJ 8, e8584, doi:10.7717/peerj.8584 (2020).

63 McMurdie, P. J. & Holmes, S. phyloseq: an R package for reproducible interactive analysis and graphics of microbiome census data. PLoS One 8, e61217, doi:10.1371/journal.pone.0061217 (2013).

64 Venables, W. N., Ripley, B. D. & Venables, W. N. Modern applied statistics with S. 4th edn, (Springer, 2002).

65 Oksanen, J. B., F. Guillaume; Friendly, Michael; Kindt, Roeland; Legendre, Pierre; McGlinn, Dan; Minchin, Peter R.; O’Hara, R. B.; Simpson, Gavin L.; Solymos, Peter; Henry M.; Stevens, H.; Szoecs, Eduard; Wagner, Helene. vegan: Community Ecology Package, <https://CRAN.R-project.org/package=vegan> (2017).

66 Love, M. I., Huber, W. & Anders, S. Moderated estimation of fold change and dispersion for RNA-seq data with DESeq2. Genome Biol 15, 550, doi:10.1186/s13059-014-0550-8 (2014).

67 Kurtz, Z. D. et al. Sparse and compositionally robust inference of microbial ecological networks. PLoS Comput Biol 11, e1004226, doi:10.1371/journal.pcbi.1004226 (2015).

68 Csardi, G. & Nepusz, T. The igraph software package for complex network research. InterJournal, Complex Systems 1695 (2006).

69 Sanchez, G. (Berkeley, 2013).

70 Team, R. C. R: A language and environment for statistical computing., <https://www.R-project.org/> (2020).

