## Supplementary Figures for "Micronutrient supplements with iron promote disruptive protozoan and fungal communities in the developing infant gut"

**Supplementary Figure 1**

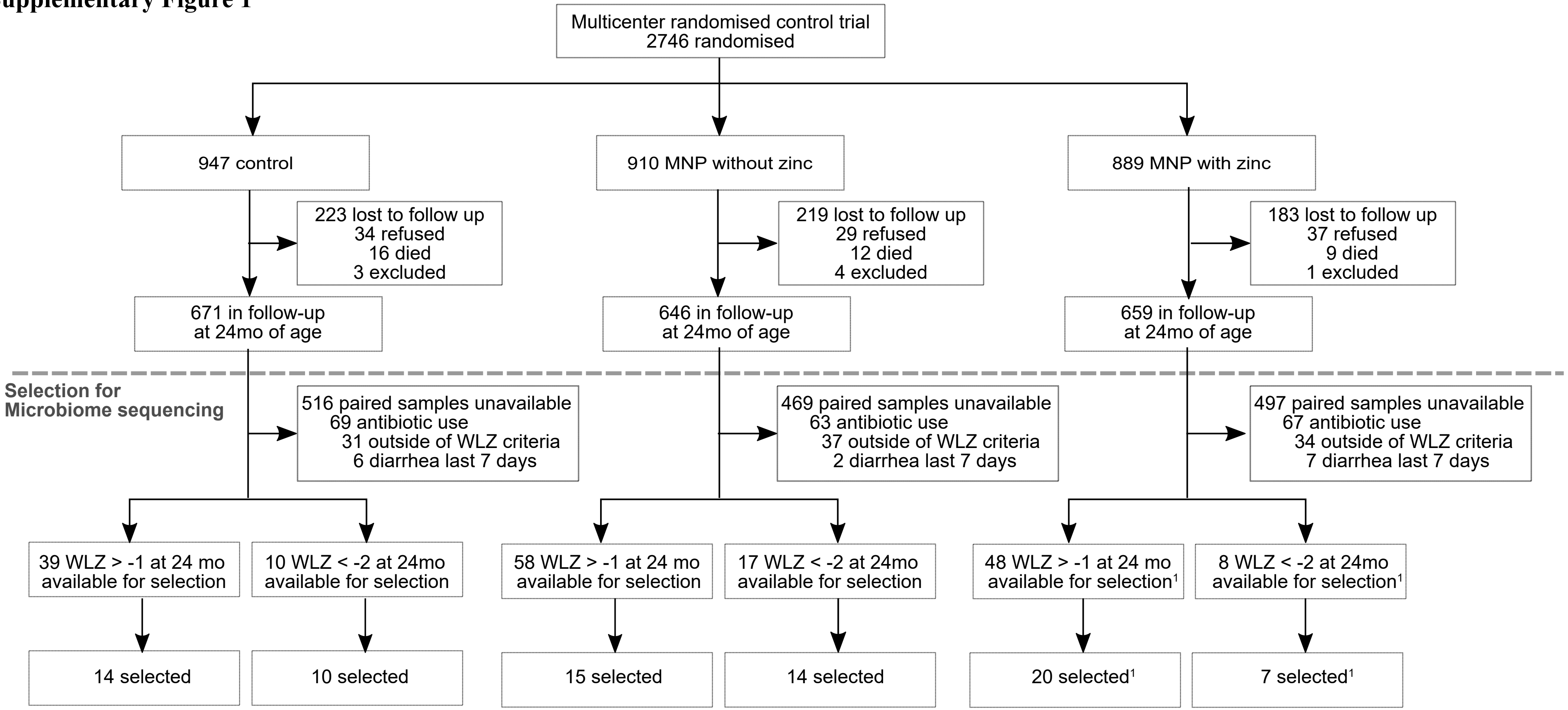


[1] Two subjects (one in the reference WLZ group and one undernourished) had, at 12 months, no diarrhea within 1 day of stool collection but reported diarrhea within 7 days prior.

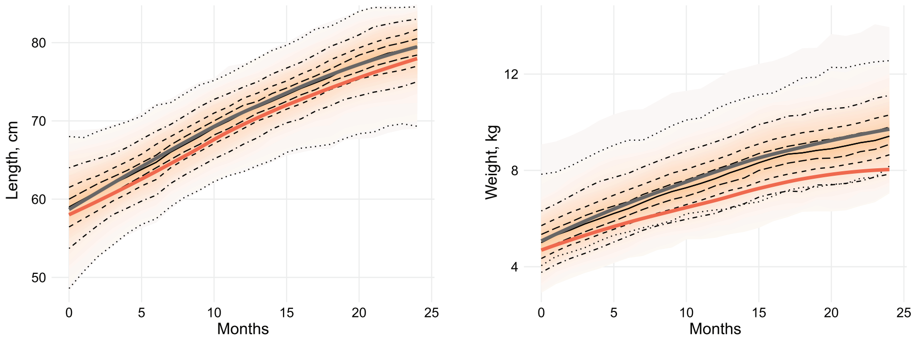


**Supplementary Figure 2.** Length (left) and weight (right) z-scores of children recruited into clinical trial NCT00705445 during the first 24 months of life. Median and quantile values are shown, with medians for participants profiled in current study indicated by red (undernourished) and black (reference WLZ) lines.

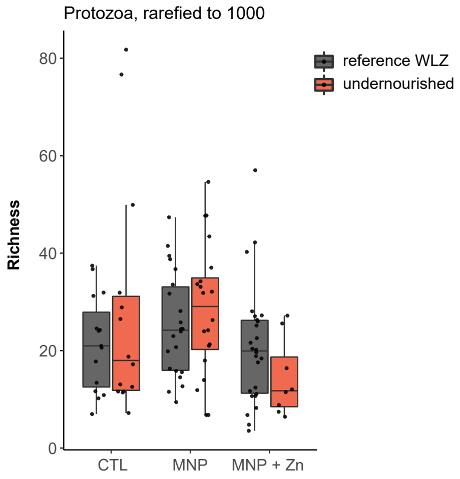


**Supplementary Figure 3.** Protozoan richness in stool samples, grouped by nutritional status and micronutrient supplementation. Samples were rarefied to 1000 reads.

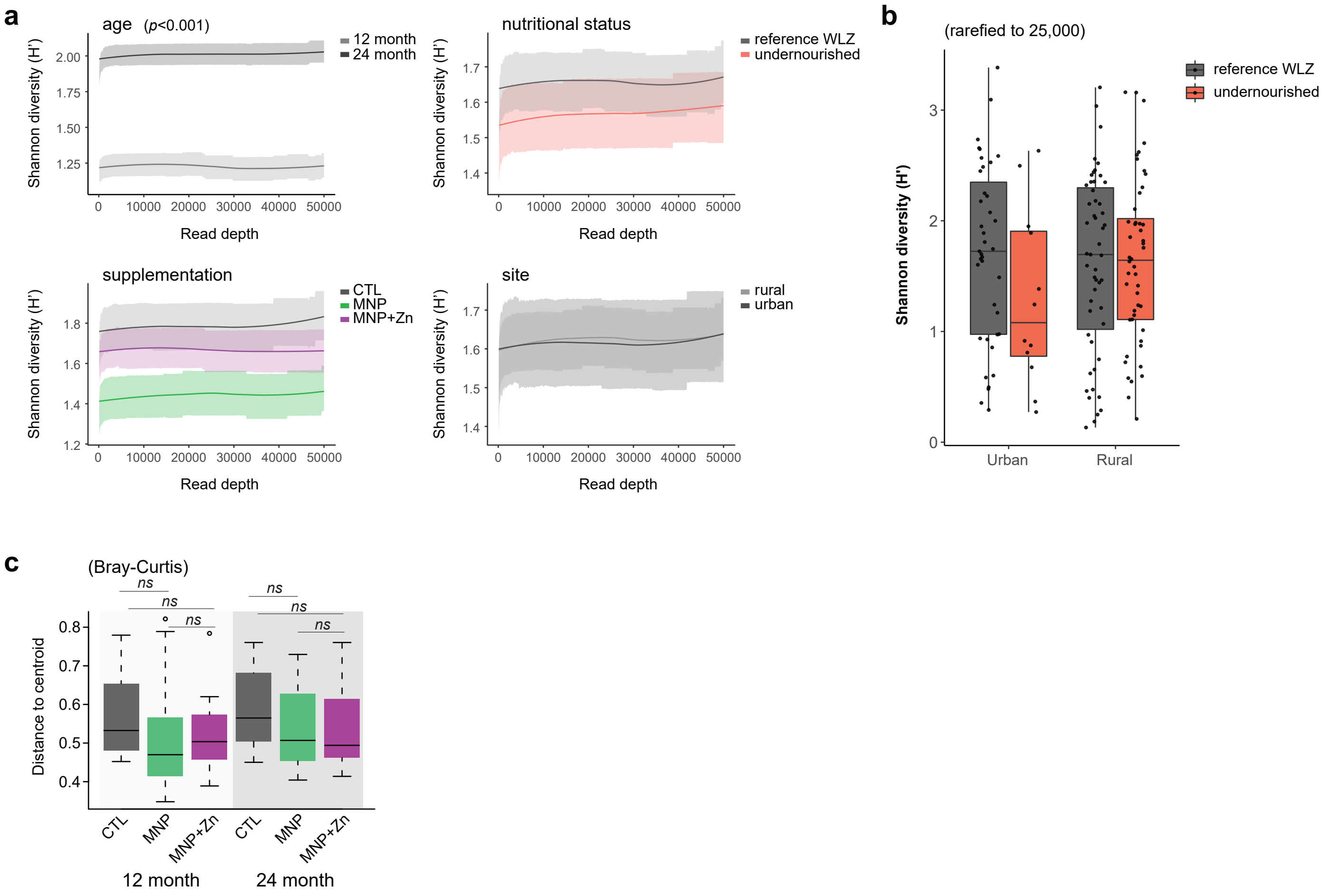


**Supplementary Figure 4.** Bacterial evenness and beta diversity. **a,** Rarefaction curves comparing species evenness (H′) by age group, micronutrient supplementation arm, nuritional status and site. Shaded regions represent standard error. Significance was tested at 25,000 read depth using a generalized linear model with stewise AIC-based model selection to identify significant variables. **b,** Boxplots comparing bacterial evenness by nutritional status and site. **c,** Variance among Bray-Curtis dissimilarities, calculated as distance to the centroid, between samples (n=150) by treatment arm and age group.

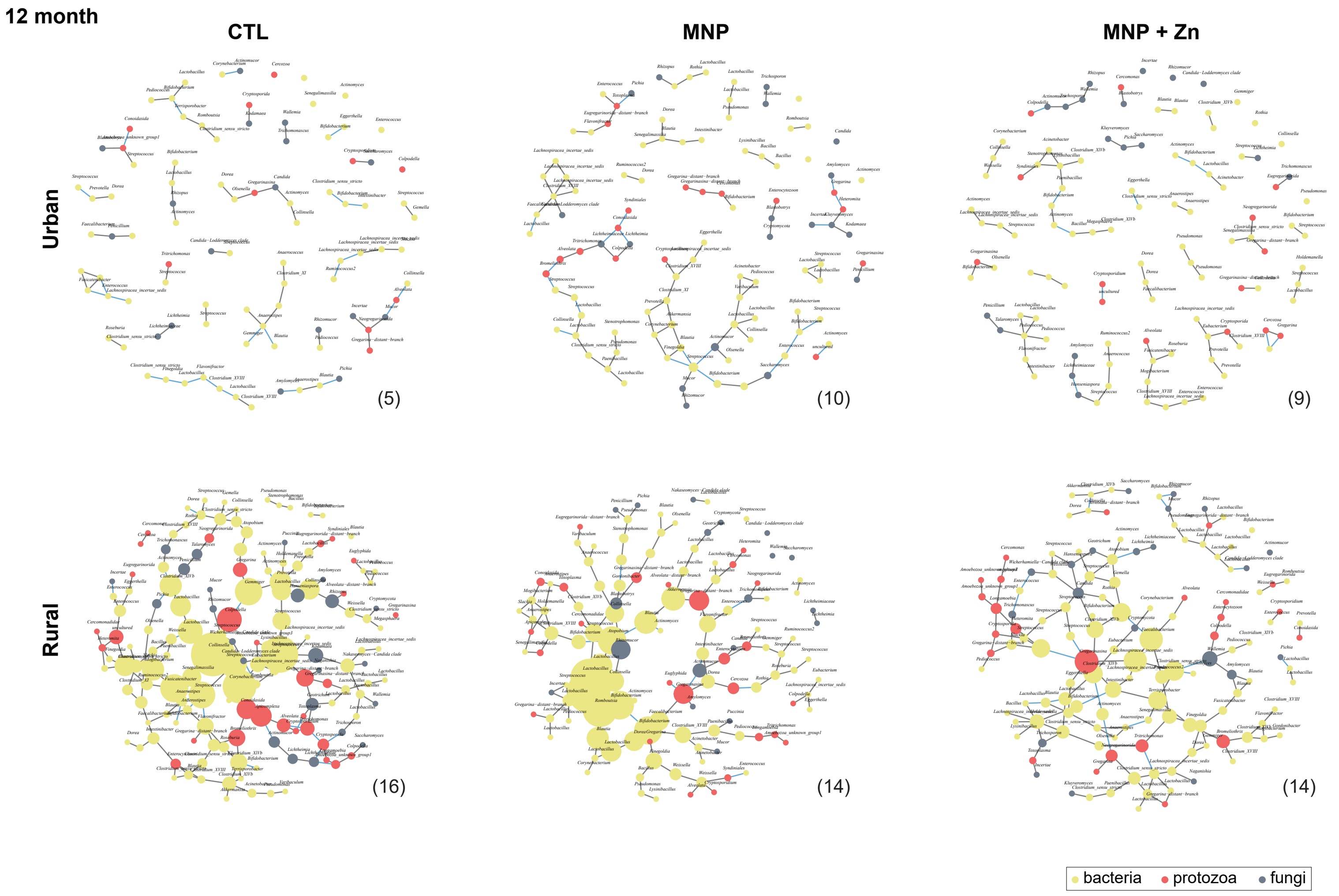


**Supplementary Figure 5.** Graphic representations of microbial networks representing predicted microbial interactions in 12 month old children, grouped by place of residence and micronutrient supplementation arm. Nodes represent bacterial OTUs (yellow) and protozoan and fungal genera (red and grey, respectively), scaled by betweenness centrality scores. Edges represent significant positive (grey) and negative (blue) correlations among microbiota. Taxa with no predicted interactions have been removed. Numbers of samples used to generate each network are indicated within brackets.
